# Circadian gene Clock regulates mitochondrial morphology and functions by posttranscriptional way

**DOI:** 10.1101/365452

**Authors:** Lirong Xu, Qianyun Cheng, Bingxuan Hua, Tingting Cai, Jiaxin Lin, Gongsheng Yuan, Zuoqin Yan, Xiaobo Li, Ning Sun, Chao Lu, Ruizhe Qian

## Abstract

Many daily activities are under the control of circadian clock, including nutrition metabolism and energy generation. Mitochondria, as the core factories of oxidizing substrates and producing ATP, undergo changes in quantity and morphology to adapt to the demand for energy. It has been demonstrated that mitochondrial gene expression, dynamics and functions are all affected by circadian clock. Here, we demonstrated that circadian gene Clock affects the number, architecture and function of mitochondria via posttranscriptional regulation of Drp1. Clock^Δ19^ leads to fragmented mitochondria accompanied with the loss of membrane potential, excessive ROS accumulation and decreased mitochondrial respiration and ATP generation. Clock^Δ19^ mice exhibit disordered lipid metabolism and evident nonalcoholic fatty liver disease (NAFLD), which are rescued by treatment with the mitochondrial fission inhibitor Mdivi-1. These results suggest a strong relationship between Clock, mitochondrial dynamics and metabolic diseases and provide a new perspective on disordered circadian clock and related diseases.

## Introduction

Circadian clock orchestrates many daily activities of living beings including sleep-wake cycle, food intake, digestion and hormone secretion. The accurate operation of circadian system relies on the transcription-translation feedback loop formed by core clock genes including Clock, Bmal1, Per, Cry, Rev-erb and downstream circadian clock-controlled genes (CCGs)(Takahashi, 2015). Among these, CLOCK, together with its heterodimer, BMAL1, are at the core, acting as positive transcription factors in the feedback loop. Unlike mice missing Bmal1, Clock^−/-^ mice appear relatively normal because of the existence of its paralog, Npas2(DeBruyne et al, 2007). Clock^Δ19^ was reported to be the most noteworthy mutation in the Clock gene, and it is accompanied by altered activity, food intake periods and apparent metabolic disorders(Turek et al, 2005), especially in terms of lipid metabolism. Clock^Δ19^ mice suffer from serious obesity and hyperlipidemia, while the Bmal1^−/-^ mice are seriously underweight(Bunger et al, 2000; Lefta et al, 2012). These differences indicate the possibility of independent regulatory roles of Clock and Bmal1, but the underlying mechanism is still under investigation.

Increasingly, studies are suggesting that many genes involved in metabolism are periodic, including genes associated with glucose and lipid metabolism(Menet et al, 2012; Neufeld-Cohen et al, 2016). Mitochondria, as the core factories oxidizing nutrient substrates and generating energy, are also under the control of circadian clock. First, quantitative proteomics and transcriptomics identified a predominant daily phase for mitochondrial mRNA and protein accumulation, including carriers involved in pyruvate and fatty acid translocation, enzymes mediating key mitochondrial metabolism and components of the mitochondrial respiration chain(Neufeld-Cohen et al, 2016). Second, circadian clock can also regulate the activities of many mitochondrial enzymes by affecting posttranslational modifications, such as the acetylation of respiration complex I, by adjusting NAD+/NADH levels and SIRTs activities(Brown, 2016; Cela et al, 2016). In addition, mitochondria are not static organelles; they experience periodic fusion and fission events(Liesa & Shirihai, 2013). It was found that Bmal1^−/-^ mice have enlarged and dysfunctional mitochondria that are less responsive to metabolic input(Jacobi et al, 2015). Additionally, mitochondrial functions and dynamics in Per1/2^−/-^ mice are disordered(Schmitt et al, 2018). However, no direct CLOCK regulation of mitochondrial dynamics has been reported.

In vivo and vitro experiments have both illustrated that altered mitochondrial dynamics give rise to metabolic block and abnormal quantity of ROS, resulting in metabolic diseases(Liesa & Shirihai, 2013; Lopez-Lluch, 2017; Yoon et al, 2006). Mitochondrial dynamics disorders are common in metabolic diseases. Mitochondria in pancreatic β-cells of patients with type 2 diabetes tend to be swollen and dysfunctional(Yoon et al, 2011). In turn, the inhibition of mitochondrial fission in β-cells results in decreased glucose-stimulated insulin secretion(Yoon et al, 2011). Similar phenomena have also been found in patients and animals with hyperglycemia(Yoon et al, 2011). Recently, Moshi Song reported a close relationship between mitochondrial dynamics and heart disease, wherein conditional deletion of Drp1 in the heart evoked dilated cardiomyopathy, while double ablation of Mfn1 and Mfn2 led to eccentric hypertrophy(Song et al, 2015). Nevertheless, there are many relationships that need to be clarified between circadian clock, mitochondrial dynamics and diseases.

In this study, we provide evidence that Clock controls the number, morphology, and functions of mitochondria by post-transcriptionally regulating mitochondrial dynamics. Disordered dynamics in Clock^Δ19^ mice led to abnormal mitochondrial architecture and mitochondrial dysfunction, which in turn led to the development of metabolic diseases. Some metabolic disorders in Clock^Δ19^ mice can be rescued by drugs that regulate mitochondrial dynamics, which suggests that Clock-controlled mitochondrial dynamics are crucial for healthy metabolism and could provide a new perspective on disordered circadian rhythms and related diseases.

## Results

### Clock^Δ19^ mice present morphological changes in their mitochondria

To balance energy supply and demand, mitochondria continually undergo changes in amount and architecture. To assess whether these mitochondrial changes were under the control of Clock, we used Clock^Δ19^ mice from the Jackson lab. First, the rhythmic activity in WT mice disappeared in Clock^Δ19^ mice and tended to be disordered (Figure S1A). In addition, Clock^Δ19^ mice gained weight faster than the same-age WT mice, with worse tolerance of insulin (Figure S1B-C). All of these suggested that Clock^Δ19^ mice experience disordered lipid and glucose metabolism. To explore mitochondrial alterations in Clock^Δ19^ mice, WT and Clock^Δ19^ livers collected at ZT0 and ZT12 were visualized by electron microscope. Mitochondria in Clock^Δ19^ mice liver were fragmented accompanied with disordered inner membrane structure (Figure 1A). Primary hepatocytes were further isolated to validate the mitochondrial changes in Clock^Δ19^ mice. First, the hepatocytes of Clock^Δ19^ mice presented an accumulation of vacuoles caused by endoplasmic reticulum swelling (Figure 1B, black arrows), which underlay cellular swelling that resulted in mitochondrial dysfunction and the lack of ATP generation. In addition, abundant lipid droplets revealed the lipid metabolism disorders in Clock^Δ19^ mice (Figure 1B, white arrows) and suggested the existence of fatty liver disease in Clock^Δ19^ mice. Moreover, the mitochondrial matrix was deeply stained, making the cristae structure appear unclear (Figure 1B, M labeled), suggesting pH changes in Clock^Δ19^ mitochondria. Furthermore, we calculated the distribution of mitochondrial surface and found most of the WT mitochondrial surface was concentrated between 0.9 and 1.1 μm^2^, while the mitochondria of Clock^Δ19^ mice were approximately 0.1-0.3 μm^2^ (Figure 1C), indicating that the mitochondria in Clock^Δ19^ mice are smaller, with a shorter diameter. We then infected the primary hepatocytes with ad-Cox8a-GFP virus. In WT livers, the mitochondria were point stained, while the point form was destroyed in the Clock^Δ19^ liver and tended to be fragmented (Figure 1D). We then used ATP6 encoded by mitochondria as a typical marker to observe mitochondrial form. The spots representing ATP6 were more specific and stronger in the WT liver compared with the diffused distribution in the Clock^Δ19^ liver (Figure 1E), demonstrating the destruction of mitochondrial architecture in Clock^Δ19^ mice.

**Figure 1.**
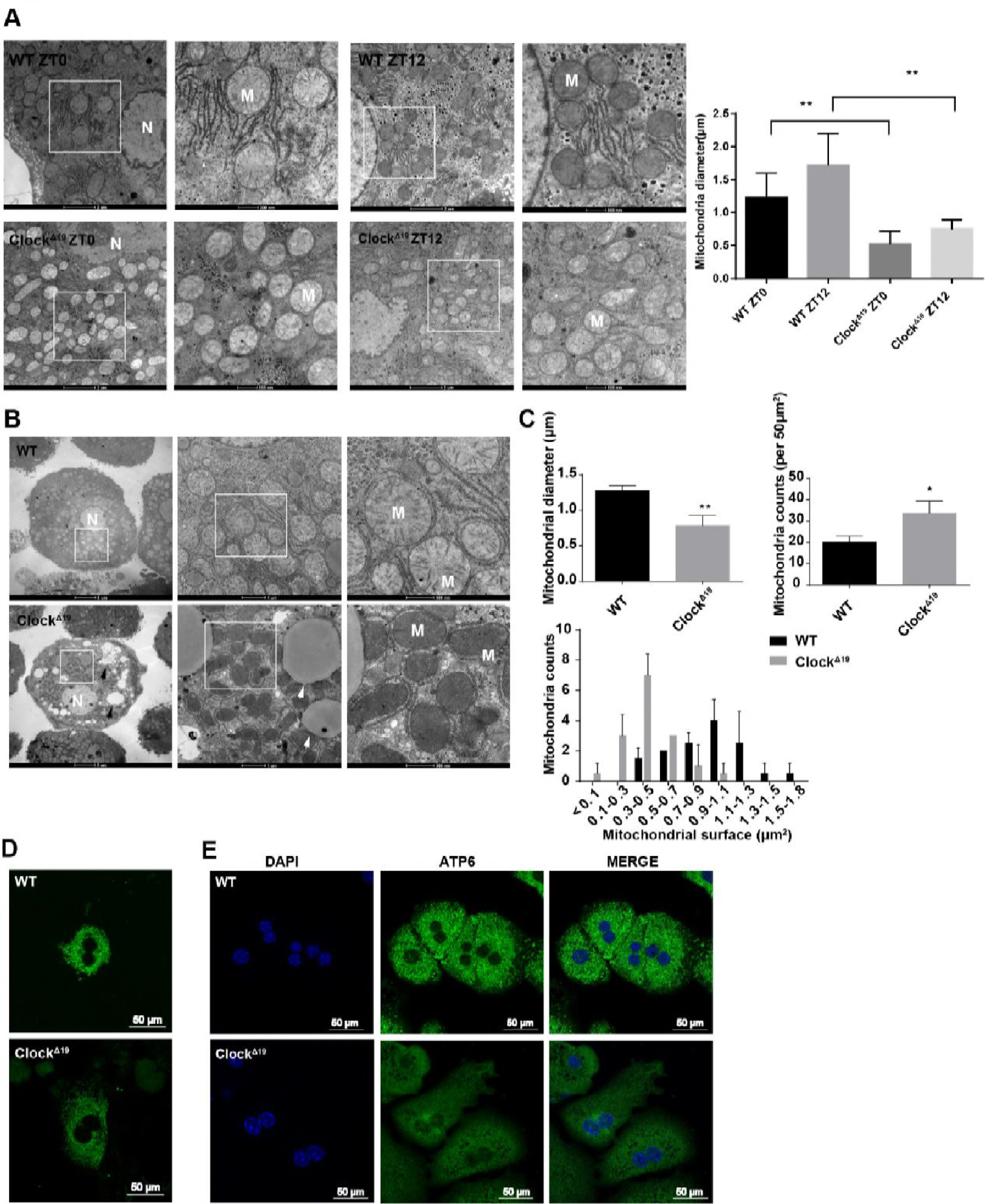
Clock^Δ19^ mice present morphological changes in mitochondria. (A) EM images of WT and Clock^Δ19^ mice livers collected at ZT0 and ZT12. Mitochondrial diameter was calculated based on the EM images. Data presented as the mean ± SEM. **p < 0.01 vs WT ZT0; **p < 0.01 vs WT ZT12 (N: Nuclear; M: Mitochondria). (B) Representative EM images of primary hepatocytes separated from WT and Clock^Δ19^ mice (Black arrows: Vacuoles; White arrows: Lipid droplets). (C) Mitochondrial diameter and size distributions calculated from EM images in 50 μm^2^. Data presented as the mean ± SEM. **p<0.01 vs WT, *p < 0.05 vs WT. (D) Confocal images of the primary hepatocellular mitochondria of WT and Clock^Δ19^ mice. The mitochondria were tagged with ad-COX8a-GFP virus. (E) Immunofluorescence images of WT and Clock^Δ19^ primary hepatocytes. Green: ATP6 stained with FITC; Blue: nucleus stained with DAPI.

### The mitochondrial morphological changes in Clock^Δ19^ mice are accompanied by dysfunction

We wondered if the abnormal mitochondria morphology could cause mitochondrial malfunction in Clock^Δ19^ mice. ROS (reactive oxygen species) were measured by fluorochrome tracing, and there was a large increase in ROS in Clock^Δ19^ primary hepatocytes compared with WT (Figure 2A and 2C). Mitochondria take in NADH produced from nutrient oxidation and generate ATP. During this procedure, the membrane potential (∆Ψ_m_) is formed, which is essential for mitochondria to provide energy. Here, JC-1 staining showed that Clock^Δ19^ was associated with a decrease in the membrane potential (Figure 2B and 2C), which also indicated that mitochondrial function is abnormal in Clock^Δ19^ hepatocytes.

**Figure 2.**
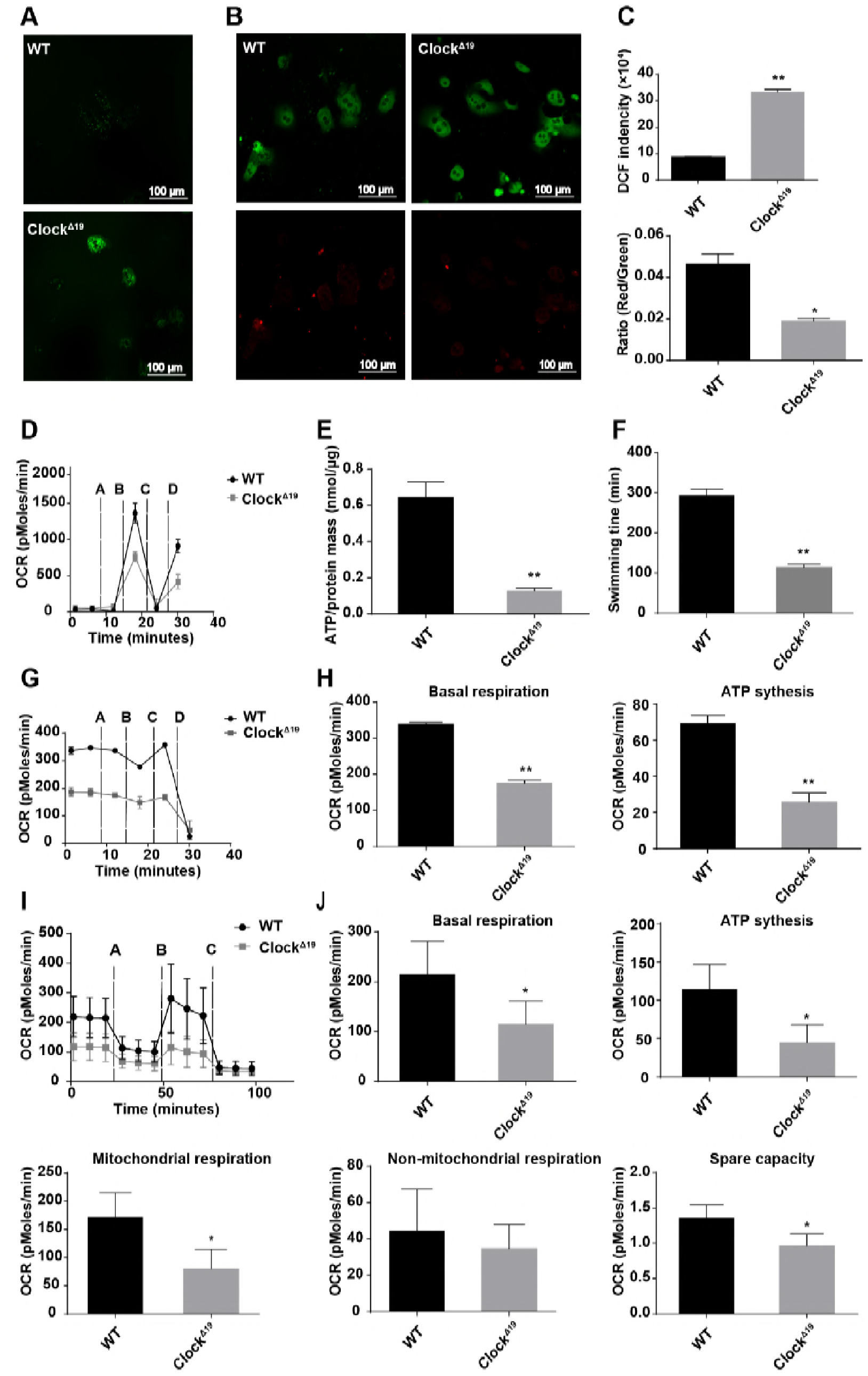
The morphological changes of Clock^Δ19^ mice are accompanied with mitochondrial respiration and ATP generation dysfunction. (A) ROS measurement images of WT and Clock^Δ19^ primary hepatocytes stained by DCFH-DA. (B) JC-1 staining of WT and Clock^Δ19^ primary hepatocytes, mitochondrial membrane potential measured by the relative fluorescence density of the red/green ratio. (C) Upper panel: Statistical histogram of ROS measurement, data presented as the mean ± SEM. **p<0.01 vs WT. Lower panel: Statistical histogram of JC-1 staining, data presented as the mean ± SEM. *p < 0.05 vs WT. (D) Electron flow assay of mitochondria separated from WT and Clock^Δ19^ livers. Injections for A-D refer to rotenone, succinate, antimycin A and ascorbate/TMPD, respectively. (E) ATP generation capability presented by the ratio of ATP concentration/total protein mass in WT and Clock^Δ19^ primary hepatocytes. Data presented as the mean ± SEM. **p<0.01 vs WT. (F) Exhaustive swimming assay conducted in WT and Clock^Δ19^ mice. Their swimming time were recorded and represent their physical power. Data presented as the mean ± SEM. **p<0.01 vs WT. (G) Coupling assay in mitochondria isolated from WT and Clock^Δ19^ livers; the injections from A to D refer to ADP, oligomycin, FCCP and antimycin A, respectively. (H) Statistical histogram of OCR represents the basal respiration and ATP synthesis determined based on the coupling assay data. Data presented as the mean ± SEM. **p<0.01 vs WT. (I) Coupling assay in primary hepatocytes. The injections from A to C refer to oligomycin, FCCP and antimycin A/rotenone, respectively. (J) Statistical histogram of OCR represents basal respiration, ATP synthesis, mitochondrial respiration, nonmitochondrial respiration and spare capacity determined based on the coupling assay data. Data are presented as the mean ± SEM. *p < 0.05 vs WT.

Mitochondria oxidize carbohydrates and lipids to generate ATP by oxidative phosphorylation. The intact mitochondria electron transport chain is critical for its ability to supply energy. It was reported that the activity of complex Ι in mitochondrial electron transport chain is Bmal1-dependent(Cela et al, 2016). To assess the impact of Clock^Δ19^ on mitochondrial complexes, we determined the electron flow of primary hepatocytes by seahorse assay. The results showed that mitochondrial respiration driven by succinate (complex II) and ascorbate/TMPD (complex IV) were clearly decreased in Clock^Δ19^ mice compared to the WT controls (Figure 2D). We then measured the ATP concentration in primary hepatocytes to evaluate the general function of mitochondria. We found that Clock^Δ19^ primary hepatocytes had a large decline in the concentration of ATP compared with WT primary hepatocytes (Figure 2E). The exhaustive swimming assay was then applied to evaluate the ATP supply and physical power in the mice. Compared to the WT mice, Clock^Δ19^ mice were more easily exhausted, revealing ATP generation and utilization dysfunctions (Figure 2F). The seahorse assay was performed to further examine mitochondrial respiration. Mitochondria isolated from Clock^Δ19^ mouse livers showed lower reactions to inhibitors (Figure 2G). Both of the OCR reflecting basal respiration and ATP synthesis were significantly reduced in Clock^Δ19^ (Figure 2H). Moreover, we measured the mitochondrial respiration of primary hepatocytes, and decreased reactions to inhibitors was also evident (Figure 2I). Both basal respiration and ATP synthesis appeared to be decreased in hepatocytes (Figure 2J). In addition, mitochondrial respiration but not nonmitochondrial respiration showed an obvious decrease in Clock^Δ19^ mice (Figure 2J). Furthermore, the decreased spare capacity in Clock^Δ19^ mice indicated a weak rapid adaption to metabolic changes (Figure 2J). Collectively, these findings revealed that the morphological changes in Clock^Δ19^ mitochondria are associated with dysfunction in mitochondrial respiration and ATP generation.

### Alterations in the expression of mitochondrial-related genes in Clock^Δ19^ mice

Unlike other organelles, mitochondria have an independent genome, although it only codes for 13 proteins and several RNAs. Mitochondrial structural components and biological functions mostly rely on the nuclear genome(Peralta et al, 2012; Scarpulla, 2008). To investigate the mechanism underlying mitochondrial dysfunction in Clock^Δ19^ mice, we examined the mRNA expression levels of several mitochondria function-related genes in livers collected every 4 h for 24 hrs. Mrps24 and Mrpl50, which are encoded by the nuclear genome and participate in mitochondrial translation, had decreased expression levels in Clock^Δ19^ mice compared to WT mice (Figure S2A). The luciferase reporter assay indicated that the transcriptions of Mrps24 and Mrpl50 are regulated by CLOCK and its heterodimer, BMAL1 (Figure S2B). On the other hand, genes encoded by the mitochondrial genome, including ND1 and ATP6, were not significantly different between WT and Clock^Δ19^ mice, except for an upregulation at ZT0 (Figure 3A). The results of the luciferase reporter assay showed that the transcription of the mitochondrial D-loop was not controlled by CLOCK or BMAL1 (Figure 3B). However, the levels of mitochondria-specific proteins including ATP5a, ATP6, ND1 and COX4 were all elevated in the Clock^Δ19^ mice total cell lysate but not in mitochondrial lysate (Figure 3C), indicating that Clock mutation does not affect the expression capacity of a single mitochondria but rather the quantity of mitochondria.

**Figure 3.**
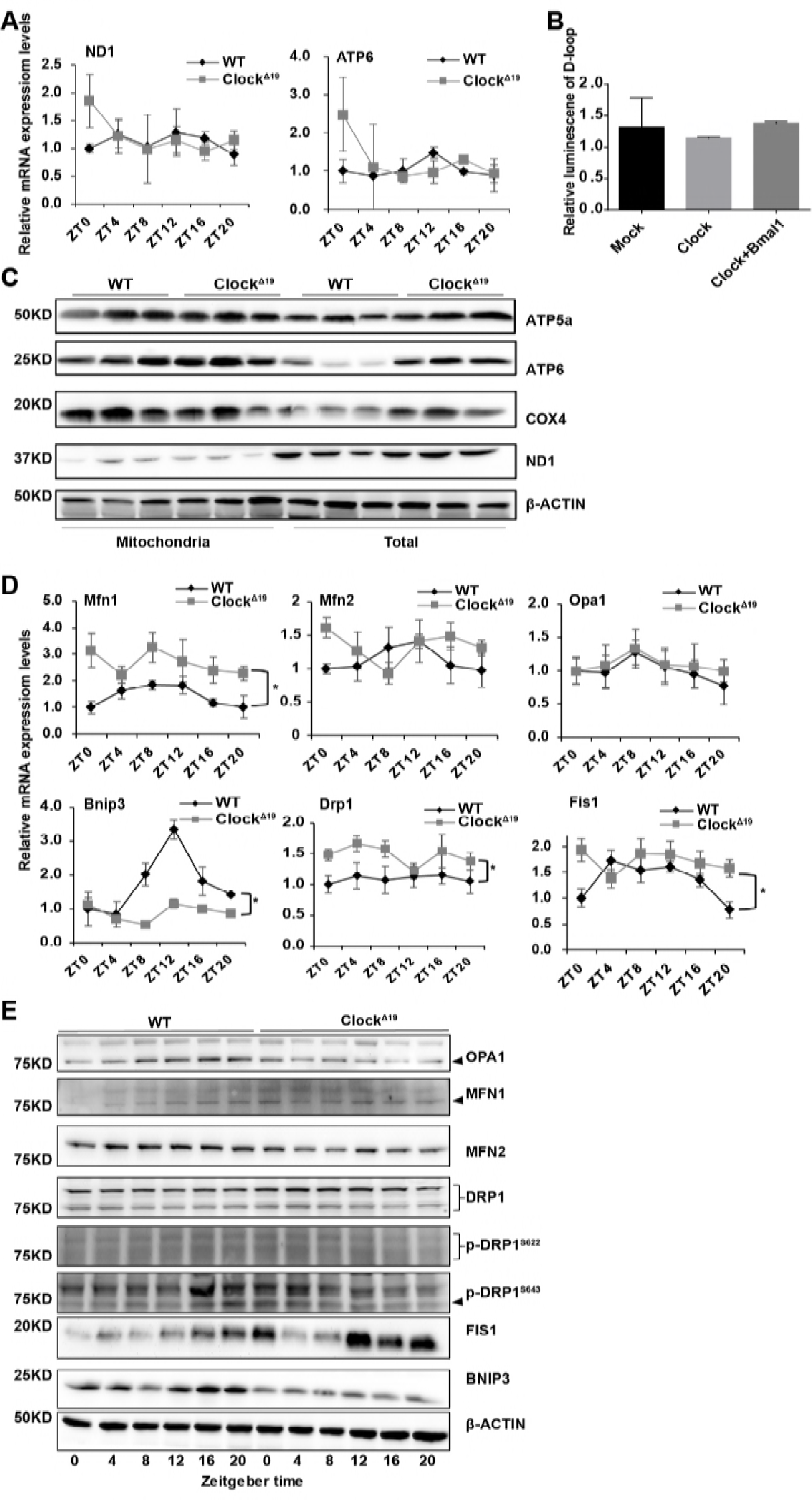
Expression of mitochondria-related genes altered in Clock^Δ19^ mice. (A) Diurnal mRNA expression of mitochondrial encoded genes in WT and Clock^Δ19^ livers measured by real-time PCR. Data presented as the mean ± SEM. (B) Luciferase reporter assay conducted in 293T cells with the mitochondrial D-loop, or together with Clock and Bmal1. Data presented as the mean ± SEM. (C) Western blot measurement of the protein expression levels of mitochondria-specific proteins in liver mitochondrial lysates and total lysates. Liver samples were collected at ZT12. (Zeitgeber time ZT0: lights on; ZT12: lights off). (D) Diurnal mRNA expression of genes involved in the regulation of mitochondrial dynamics in WT and Clock^Δ19^ livers measured by real-time PCR. Data presented as the mean ± SEM.*p < 0.05 vs WT. (E) Western blot analysis of liver mitochondrial dynamics-related proteins. The liver tissues were collected every 4 h for 24 hrs.

We next sought to determine the molecular mechanisms underlying excessive mitochondrial fission in Clock^Δ19^ mice. We used RT-PCR to detect the mRNA expression levels of genes involved in mitochondrial dynamics. The results showed that, except for a slight elevation of Mfn1 in Clock^Δ19^ mice, there were no significant differences in other fusion-related genes between Clock^Δ19^ and WT mice. Meanwhile, Drp1 and Fis1, the primary mitochondrial fission genes had increased expression levels in Clock^Δ19^ mouse livers compared with WT mouse livers. Clearly, Bnip3, a main mitophagy gene, which was reported to be rhythmically expressed, decreased and failed to cycle in Clock^Δ19^ mouse livers (Figure 3D). Moreover, the expression levels of fusion proteins (Opa1, Mfn1 and Mfn2) were reduced in Clock^Δ19^ mouse livers, although not as significantly as Bnip3 (Figure 3E). In contrast, the expression levels of fission proteins (Drp1 and Fis1) were increased in Clock^Δ19^ mice compared to WT mice (Figure 3E). Phosphorylation of DRP1 is important for its GTPase activity, which then influences mitochondrial fission. Phosphorylation at s622 (s616 in humans) increases fission, while phosphorylation at s643 (s637 in humans) plays a negative role in mitochondrial fragmentation(Chang & Blackstone, 2007; Chang & Blackstone, 2010). However, in our research, we did not find any significant differences in DRP1 phosphorylation between WT and Clock^Δ19^ mice (Figure 3E).

### Excessive mitochondrial fission in Clock^Δ19^ mice is due to posttranscriptional regulation of Drp1 by CLOCK

To further investigate the changes in protein expression, we used immunofluorescence to study the alterations in the expression of FIS1 and DRP1; the staining intensity showed similar trends with the Western blot (Figure 4A). By analyzing ChIP-Seq data, we found no significant accumulation of CLOCK on the promoter region of Drp1(Annayev et al, 2014), indicating that Drp1 is probably regulated by Clock in a posttranscriptional manner. Moreover, the increase of DRP1 in Clock^Δ19^ mice was eliminated by the translational inhibitor cycloheximide but not by actinomycin D, which is a transcriptional inhibitor (Figure 4B), suggesting that the accumulation of DRP1 in Clock^Δ19^ mice is not attributable to slower protein degradation but rather to more protein generation or less mRNA degradation. To compare the degradation rate of Drp1 mRNA in WT and Clock^Δ19^ mice, we measured the changes in relative mRNA expression levels over time after the addition of actinomycin D. Slower degradation of Drp1 in Clock^Δ19^ mice verified the possibility of posttranscriptional regulation of Drp1 by Clock (Figure 4C).

**Figure 4.**
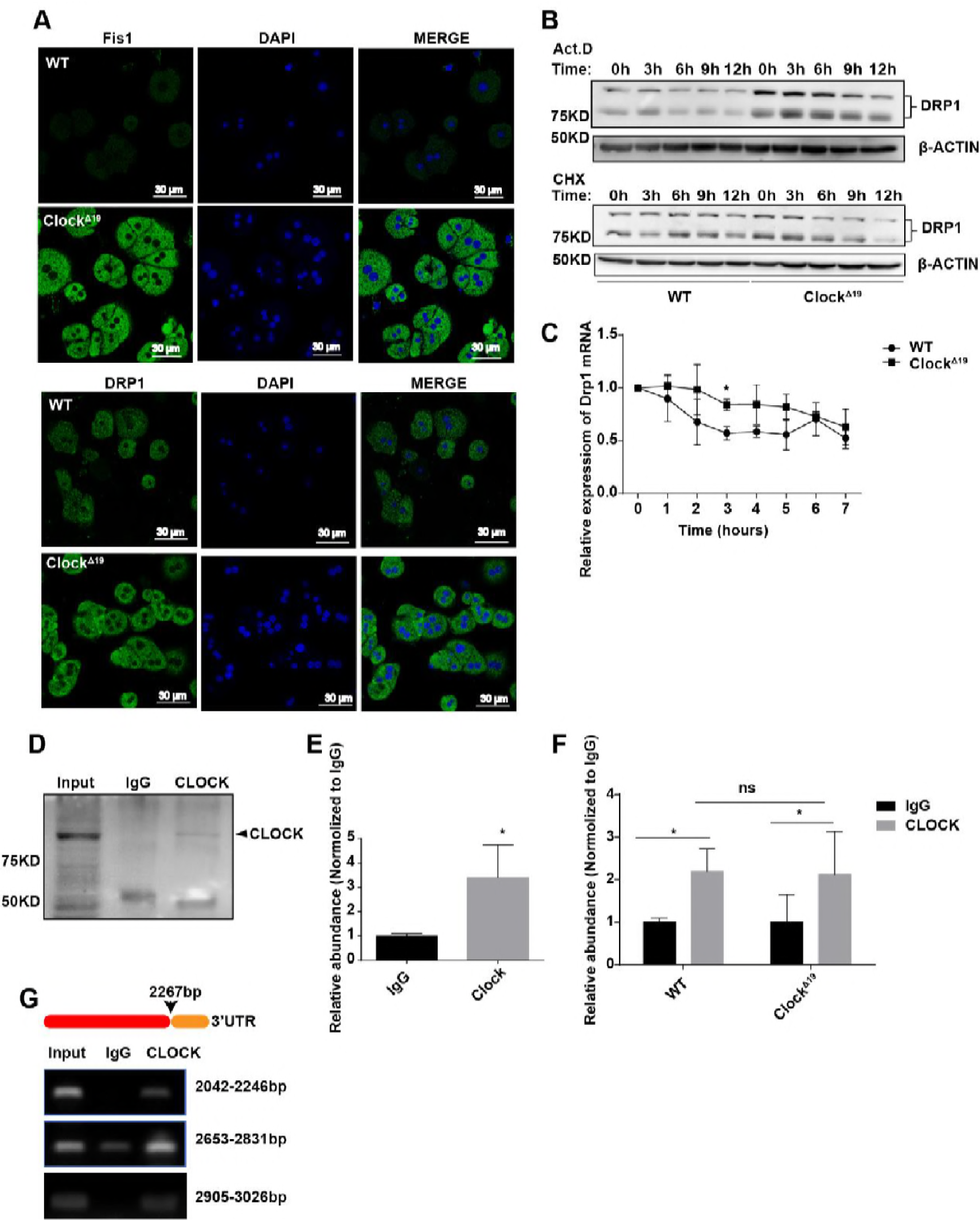
Excessive mitochondrial fission in Clock^Δ19^ is due to posttranscriptional regulation of Drp1 by CLOCK. (A) Immunofluorescence images of WT and Clock^Δ19^ primary hepatocytes. Green: FIS1/DRP1 stained with FITC; Blue: nucleus stained with DAPI. (B) Protein expression of DRP1 in WT and Clock^Δ19^ hepatocytes after treatment with 5 μg/ml Act.D or 50 μg/ml CHX for 0, 3, 6, 9 and 12 hours. (C)Relative mRNA expression level of Drp1 in WT and Clock^Δ19^ hepatocytes after treatment with 5 μg/ml Act.D for 0, 1, 2, 3, 4, 5, 6 and 7 hours. Data presented as the mean ± SEM. *p<0.05 vs WT. (D) Western blot detection of CLOCK after the RIP assay. (E) RIP analysis of AML12 lysate with IgG or anti-CLOCK antibodies. Precipitated mRNA was detected by RT-PCR. Data presented as the mean ± SEM. *p<0.05 vs IgG. (F) RIP analysis of WT and Clock^Δ19^ liver lysate with IgG or anti-CLOCK antibodies. Precipitated mRNA was detected by RT-PCR. Data presented as the mean ± SEM. *p<0.05 vs IgG. (G) CLOCK binding sites on Drp1 mRNA checked by PCR with specific primers and DNA gel analysis.

It has been reported that CLOCK can act as an mRNA splicer together with other splicing factors(Yang et al, 2018). We performed RIP assay to detect whether CLOCK can bind to Drp1 mRNA and further affect its stability. The RIP results showed that there was a significant accumulation of CLOCK on Drp1 mRNA relative to the negative control IgG in the AML12 hepatocyte cell line (Figure 4D-E). Moreover, the binding was still significant in Clock^Δ19^ mouse livers (Figure 4F), suggesting that the 19^th^ exon of Clock is not critical for mRNA binding but may play a role in interacting with other mRNA binding factors or affect its degradation-mediating activity. Moreover, by predicting the Drp1 mRNA-protein binding sites, we designed several primers to search for the specific CLOCK binding sites. It was shown that there were significant accumulations of CLOCK at 2042-2246 bp, 2653-2831 bp and 2905-3026 bp of Drp1 mRNA (Figure 4G), which are all near or in its 3’UTR region. In brief, these findings suggest that CLOCK can bind to Drp1 mRNA 3’ UTR region and further affect its stability. Less Drp1 degradation in Clock^Δ19^ mice led to abnormal mitochondrial dynamics and dysfunction.

### Mdivi-1 rescued mitochondrial morphological and functional changes

Considering the excessive fission and the severe dysfunction of the mitochondria of Clock^Δ19^ mice, we wondered whether inhibiting fission would rescue these mitochondrial changes. Mdivi-1 is an efficient mitochondrial fission repressor that works by inhibiting the GTPase activity of Drp1. Incubation with Mdivi-1 for 12 or 24 hours both strengthened the fluorescence intensity and recovered the excessively fragmented mitochondria to its granular or even tubular structures in Clock^Δ19^ hepatocytes but not in WT hepatocytes (Figure 5A), which indicates that excessive fission in Clock^Δ19^ is mainly due to abnormal Drp1 regulation. Moreover, the addition of Mdivi-1 substantially reduced ROS accumulation (Figure 5B) and restored the decreased membrane potential in Clock^Δ19^ mice comparing with the control (Figure 5C). Measurement of ATP showed an obvious restoration of mitochondrial ATP generation by Mdivi-1 in Clock^Δ19^ primary hepatocytes but not in the WT (Figure 5D).

**Figure 5.**
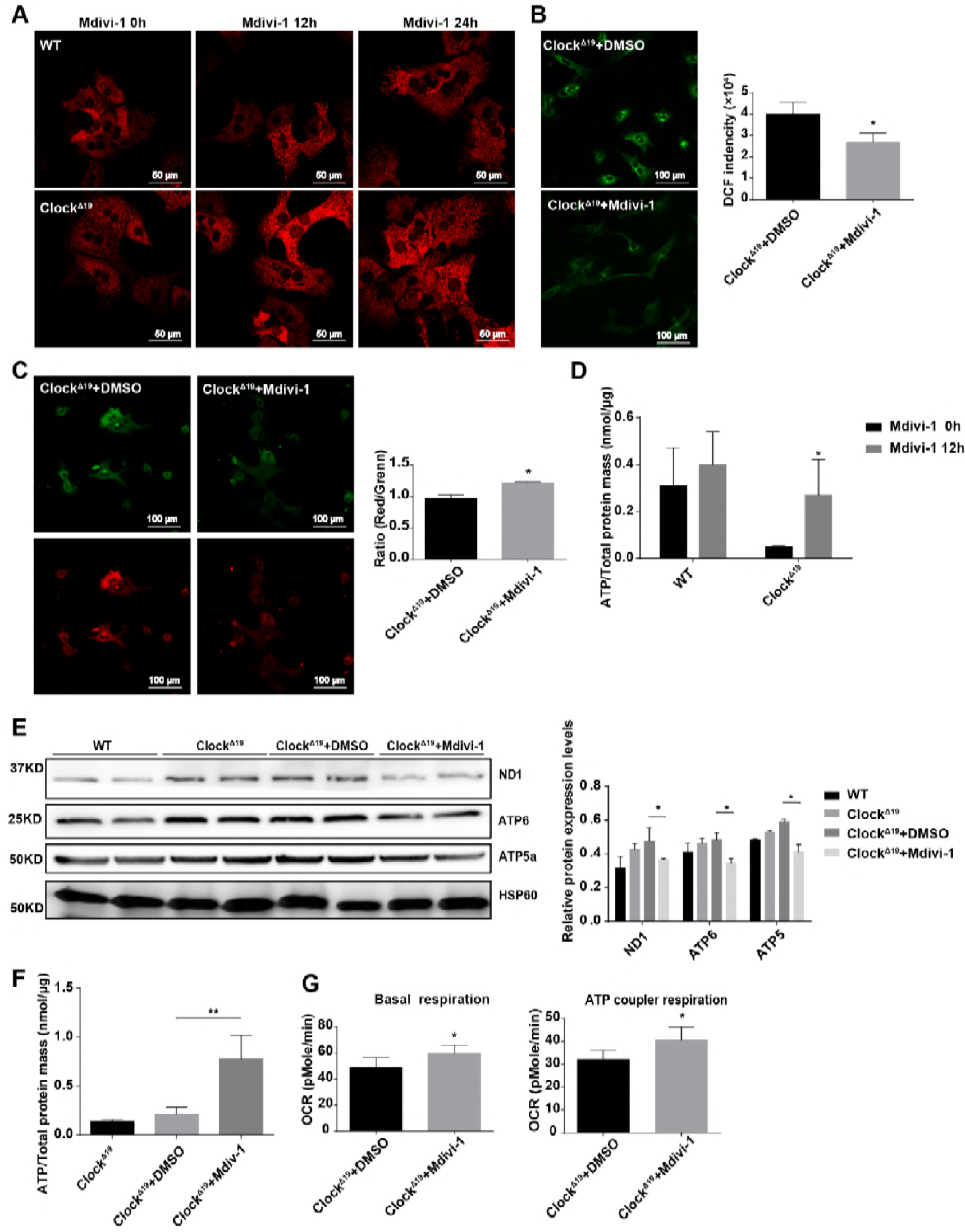
Mdivi-1 rescued mitochondrial morphology and functional changes. (A) Representative confocal images of the mitochondrial network in WT and Clock^Δ19^ primary hepatocytes with the addition of Mdivi-1 for 0, 12 and 24 hours. Mitochondria were tagged with ad-COX8a-RFP virus. (B) Total ROS production of Clock^Δ19^ primary hepatocytes with addition of DMSO or Mdivi-1, as measured by DCFH-DA staining. Right panel: statistical histogram of fluorescence intensity. Data presented as the mean ± SEM. *p < 0.05 vs Clock^Δ19^+DMSO. (C) Mitochondrial membrane potential of Clock^Δ19^ primary hepatocytes with the addition of DMSO or Mdivi-1 stained with JC-1. Right panel: statistical histogram of red/green ratio. Data presented as the mean ± SEM. *p < 0.05 vs Clock^Δ19^+DMSO. (D) ATP concentrations in WT and Clock^Δ19^ primary hepatocytes treated with Mdivi-1 for 0 and 12 hours. Data presented as the mean ± SEM. *p < 0.05 vs Clock^Δ19^+Mdivi-1-0 h. (E) Western blotting of the mitochondrial proteins of WT and Clock^Δ19^ mice livers injected with DMSO and Mdivi-1. Right panel: statistical histogram of relative protein expression levels. Data presented as the mean ± SEM. *p < 0.05 vs Clock^Δ19^+DMSO. (F) ATP production capability measurement of Clock^Δ19^ primary hepatocytes after treatment with DMSO or Mdivi-1. Data presented as the mean ± SEM. **p < 0.01 vs Clock^Δ19^+DMSO. (G) Statistical histogram of OCR represents basal respiration and ATP synthesis based on the coupling assay of the treated mice. Data presented as the mean ± SEM. *p < 0.05 vs Clock^Δ19^+DMSO.

Considering the favorable effect of Mdivi-1 on hepatocytes, we then explored its effect on mouse mitochondrial function and related metabolism by intraperitoneal injection. The obvious downregulation of the main mitochondrial proteins (ATP5a, ND1, ATP6) in liver tissue demonstrated the inhibition of mitochondrial fission by Mdivi-1 compared with DMSO (Figure 5E). Moreover, decreased ROS accumulation and increased mitochondrial membrane potential both indicated a positive effect of Mdivi-1 on Clock^Δ19^ mouse mitochondrial function by injecting (Figure S3A-B). In addition, the evident increase in the concentration of ATP and the recovery of mitochondrial respiration also serve as direct evidence for the therapeutic effect of Mdivi-1 on mitochondrial dysfunction (Figure 5F-G).

### Intraperitoneal injection of Mdivi-1 recovered hyperlipoidemia and nonalcoholic fatty liver disease in Clock^Δ19^ mice

After two weeks of injection, serum samples were collected from the treated and control mice, and IPGTT, blood glucose test and lipid test were conducted. There was no significant difference in serum glucose levels between groups, but there was some recovery on the IPTGG in the Mdivi-1-treated group (Figure 6A-B). As shown in Figure 6B, Mdivi-1-treated mice exhibited stronger tolerance than the untreated and DMSO-treated Clock^Δ19^ mice. Their blood glucose levels were slightly elevated at 15 min after glucose injection, when the glucose levels in the control groups reached their peak. These findings suggested that Mdivi-1 treatment may directly or indirectly affect glucose metabolism. In addition, the effect of Mdivi-1 on serum lipid levels is more complicated; there was a slight downregulation of triglyceride (TG) and density lipoprotein cholesterol (LDL-C) levels in the Mdivi-1-treated group compared with the DMSO control group (Figure 6C). The effect of Mdivi-1 was most evident in increasing the level of HDL-C, implying a protective function of Mdivi-1 against hyperlipoidemia and related diseases (Figure 6C). However, there was no significant glucose or lipid content improvement in WT mice receiving Mdivi-1, but a slight increase in TG (Figure S4A-B), indicating that Mdivi-1 has no or possibly even a negative effect on well-balanced mitochondria. We then used oil-red staining to confirm the rescue effect of Mdivi-1 on fatty liver diseases. Frozen section of livers showed an excellent therapeutic effect of Mdivi-1 in Clock^Δ19^ mouse livers, which suffered from serious lipid droplets deposition (Figure 6D); however, there was no significant difference seen in WT livers.

**Figure 6.**
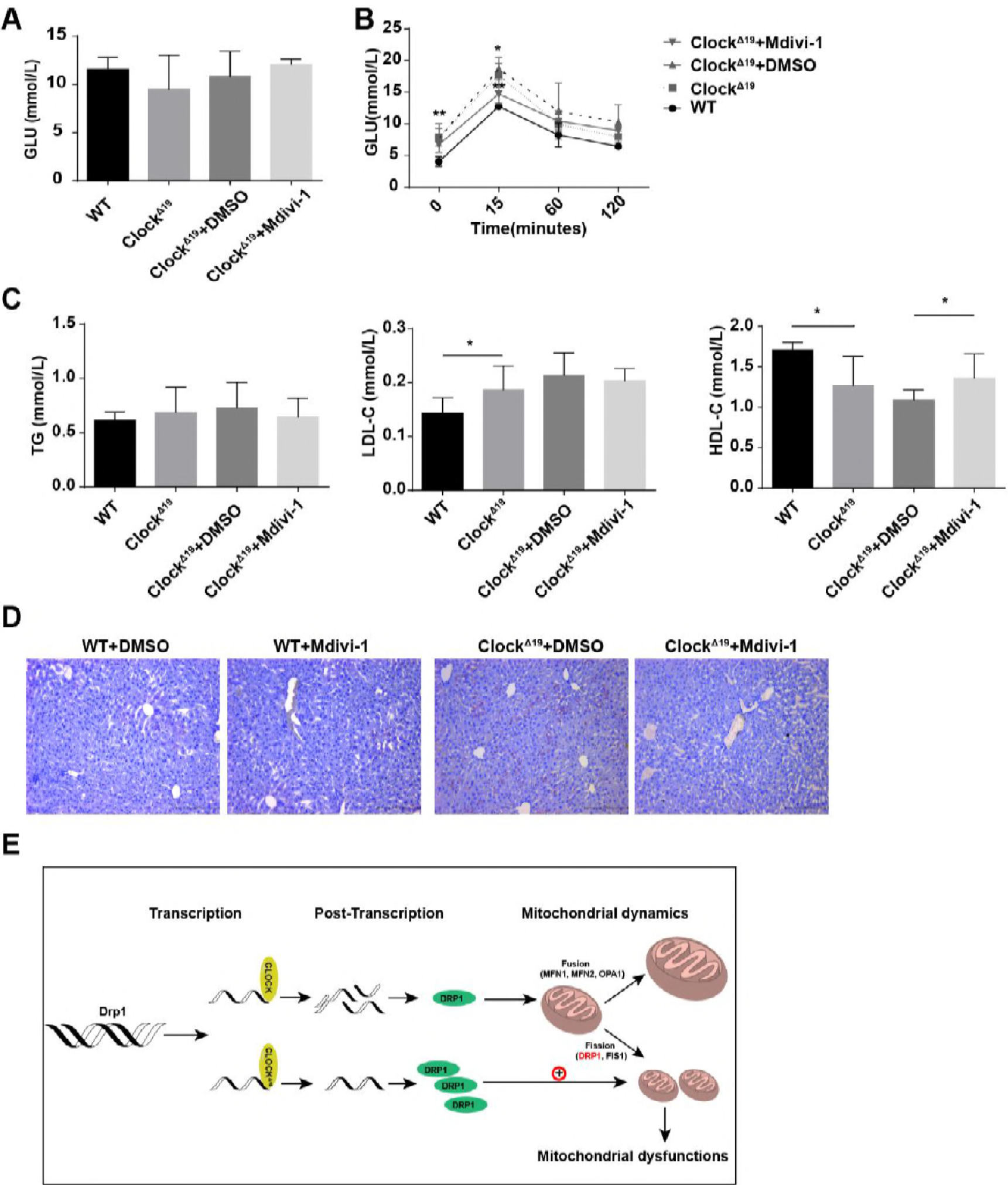
Intraperitoneal injection of Mdivi-1 rescued hyperlipidemia and nonalcoholic fatty liver disease in Clock^Δ19^ mice. (A) Fasting blood glucose levels in Clock^Δ19^ mice measured after overnight fasting (n=7). Data presented as the mean ± SEM. (B) IPGTT of WT and Clock^Δ19^ mice measured after overnight fasting (n=7). Data presented as the mean ± SEM. *p < 0.05, ClockΔ19+Mdivi-1 vs Clock^Δ19^+DMSO; **p < 0.01 Clock^Δ19^ vs WT. (C)TG, LDL-C and HDL-C in WT, Clock^Δ19^ and treated mice (n=7). Data presented as the mean ± SEM. *p < 0.05, Clock^Δ19^ vs WT, Clock^Δ19^+Mdivi-1 vs Clock^Δ19^+DMSO. (D) Oil-red staining of WT and Clock^Δ19^ liver frozen sections to measure lipid deposition. (E) A schematic diagram illustrating the role of CLOCK in mitochondrial dynamics.

## Discussion

In our study, we demonstrated that circadian gene Clock plays an important role in mitochondrial morphology and function by posttranscriptional regulation of Drp1. Furthermore, mitochondrial architecture, membrane potential, ROS production and respiration tend to be abnormal in Clock^Δ19^ mice due to the excessive mitochondrial fragmentation.

Circadian clock plays critical roles in maintaining mitochondrial functions. Either the accumulation of many mitochondrial proteins or the related functional features such as oxygen consumption, membrane potential, ATP generation and mitochondrial respiration are all rhythmic(Schmitt et al, 2018). The mitochondrial circadian oscillations are abolished in Bmal1^−/-^ and Per1/2^−/-^ mice (11,12). In our study, we provide evidence that the mutation of Clock also results in disordered mitochondrial functions. Decreased Δφm, abundant ROS accumulation, and decreased ATP concentration and mitochondrial respiration in Clock^Δ19^ verified the role of Clock in mitochondria. It was reported that the activity of mitochondrial respiration complex Ι oscillates and peaks at ZT12 mainly because of the Nampt-NAD-SIRTs pathway(Cela et al, 2016). Here, we used the seahorse assay and determined that the activities of mitochondrial respiration complexes II and IV are both decreased in Clock mutant mice, which indicates that there are other mechanisms of circadian clock regulation of mitochondrial respiration. Indeed, it has been reported that several mitochondrial complex components including NDUFA2, NDUFB5, and NDUFC1 in complex Ι; COX4I1, COX6A1, and COX7A2 in complex IV; and ATP5G2 and ATP5L in complex Ⅴ are all transcriptionally regulated by circadian clock elements(Schmitt et al, 2018). In addition, except for the regulation of acetylation by SIRTs, recent global acetylome analyses have demonstrated that the acetylation of many mitochondrial proteins is rhythmic. Moreover, some acetylation is altered or abolished in Clock^−/-^ mice(Masri et al, 2013), indicating that CLOCK plays an independent role in mitochondrial regulation as an acetyltransferase.

Recent studies have demonstrated that circadian factors Bmal1 and Per1/2 both play roles in the regulation of mitochondrial morphology and affect the related functions. Bmal1^−/-^ mitochondria tend to be enlarged and accompanied by elevated ROS levels and mitochondrial dysfunction(Jacobi et al, 2015). Per1/2 is involved in the regulation of mitochondrial dynamics by affecting the rhythmic activity of DRP1(Schmitt et al, 2018). In our study, Clock^Δ19^ mice mitochondria were fragmented with smaller diameters and greater quantity in the same area when compared with the WT. It has been reported that mitochondrial dynamics are regulated by circadian genes, mostly through their transcriptional regulation of mitochondrial dynamics genes or energy sensors(Jacobi et al, 2015; Schmitt et al, 2018). In this study, we demonstrated that the core mitochondrial fission mediator, DRP1, is under direct posttranscriptional regulation of CLOCK. CLOCK facilitates the degradation of Drp1 mRNA by binding to its 3’UTR region. In Clock^Δ19^ mice, decreased DRP1 degradation leads to DRP1 accumulation and then induces abnormal mitochondrial fission. The fact that CLOCK-Drp1 mRNA binding is still evident in Clock^Δ19^ mice suggests that the 19^th^ exon of CLOCK is not involved in its binding to mRNA but rather in some other processes. It is well known that Clock^Δ19^ nearly loses its transcriptional activity, although its interaction with BMAL1 is still evident(Gekakis et al, 1998). The underlying mechanism is still under investigation; we speculate that the 19^th^ exon of CLOCK may play a similar role in the regulation of CLOCK’s mRNA regulating activity. In addition, yeast two-hybrid of CLOCK (1-580) and the 19^th^ exon deletion mutation of CLOCK proved that CLOCK can interact with other proteins except for BMAL1 and that the 19^th^ exon of CLOCK is important in the CLOCK-CIPC complex formation(Gekakis et al, 1998; Hou et al, 2017).

In our study, we determined the mRNA degradation-mediating function of CLOCK for the first time, which is one of the critical posttranscriptional regulation processes. Indeed, except for their transcriptional regulatory roles, circadian clock factors can also engage in posttranscriptional regulation to generate circadian biological processes. As soon as transcription is initiated, mRNA modification starts; alternative splicing(McGlincy et al, 2012; Yang et al, 2018) is regulated by circadian clock, and disruption of splicing would also lead to disordered circadian rhythm by influencing mRNA translocation or stability(Green, 2017). After being synthesized, mRNAs translocate into the cytosol and are translated into proteins. It has been reported that mRNA translocation(Chen et al, 2008) and translation are both circadian-regulated processes(Green, 2017) and that the core circadian factor, BMAL1, also participates in translation by interacting with several transcriptional factors, including transcription initiation factors, elongation factors and ribosome subunits(Lipton et al, 2015). Moreover, mRNA and protein stability(Wang et al, 2018) and posttranslational modifications are also regulated by circadian clock. For example, CLOCK is not only a transcription factor but also an acetyltransferase(Doi et al, 2006). Except for the previously reported substrates such as histone and BMAL1(Doi et al, 2006; Hirayama et al, 2007), more substrates have been found, such as ASS1(Lin et al, 2017), which is an important enzyme in the urea cycle. The mRNA degradation-mediating function of CLOCK uncovered in our study supplements the general understanding of the posttranscriptional regulatory functions of the circadian clock.

Due to its vital role in the regulation of processes essential for daily life, disordered circadian rhythm and mutant circadian genes both lead to serious diseases. Atherosclerosis, aging, prostate cancer and metabolic disease are all circadian-related diseases. Abnormal activity, food intake and metabolic rates were revealed when Clock^Δ19^ mutation was first identified. As a result, Clock mutant mice gained weight faster than WT mice when fed regular or high-fat diets (Turek et al, 2005) (Adamovich et al, 2014; Aviram et al, 2016). In our study, Clock^Δ19^ mice had evident hyperlipidemia and NAFLD, suggesting they suffered from lipid metabolism disorders. It has been reported that mitochondria tend to be round and swollen in NAFLD patients, and excess ROS production is thought to be the main cause of the pathological progression. The accumulation of ROS may be attributed to excessive mitochondria fission resulted from fat accumulation(Galloway & Yoon, 2013). Abundant ROS accumulation, excessive fission and the rescuing effect of Mdivi-1 on NAFLD in Clock^Δ19^ mice verified that abnormal mitochondrial dynamics in Clock^Δ19^ mice is a cause of hyperlipoidemia and NAFLD. In vitro addition of Mdivi-1 caused a rapid reversible dose-dependent formation of net-like mitochondria. In vivo application of Mdivi-1 has been shown to protect cardiomyocytes and kidney, neuro and retinal cells following ischemia/reperfusion (Brooks et al, 2009; Ong et al, 2010), functioning as a protector against acute injury. In addition, it also plays a long-term therapeutic role in heart failure(Givvimani et al, 2012) and cardiac hypertrophy. In our study, in addition to its roles in decreasing ROS production and recovering mitochondrial membrane potential and respiration, we provide a new role for Mdivi-1 in curing hyperlipoidemia and fatty liver disease. In addition to the direct regulation of lipid metabolism-related genes by Clock, here we offer a new perspective wherein Clock affects lipid metabolism and related diseases by regulating mitochondrial dynamics.

In brief, we propose that Clock plays an important role in the regulation of mitochondrial dynamics and related functions by the posttranscriptional regulation of Drp1 (Figure 6E). The findings in our study provide a new perspective on the circadian clock, mitochondria and metabolism, which might contribute to the understanding and generate new ideas for clinical application.

## Materials and methods

### Animal studies

The Clock^Δ19^ mice were purchased from the Jackson Laboratory and had been bred in the Model Animal Research Center of Nanjing University. Same aged C57BL/6J mice were also purchased from Model Animal Research Center of Nanjing University. Mice were fed a chow diet and raised in a clean room with 12 h light and 12 h dark cycles (Lights on at 8:00am and lights off at 20:00pm). All animal experiments were conducted strictly in accordance with the National Institutes of Health Guide for the Care and Use of Laboratory Animals and were approved by the Animal Care and Use Committee of Shanghai Medical College, Fudan University.

For tissue collection, mice were sacrificed at the age of 8-12 weeks. Their tissues were harvested every 4 h for 24 hour and then stored at −80℃ or in 4% paraformaldehyde.

For metabolic index detection, mice at the age of 10 weeks with an approximate weight of 23–25 grams were divided into blank control group, vehicle control and Mdivi1-treated group. After 2 weeks of treatment, the animals were first tested with the intraperitoneal glucose tolerance test (IPGTT) and then euthanized. Their sera were collected, and the levels of blood glucose and lipids were measured. For IPGTT measurement, mice were fasted overnight for approximately 14 hours with free access to water. After the measurement of their fasting blood glucose levels, the mice were intraperitoneally injected with 20% glucose dissolved in 0.9% NaCl (1 g/kg). The blood glucose levels were then measured at 15 min, 60 min and 120 min after injection. Blood samples were obtained from the tail veins, the glucose levels were measured by glucose meter (Abbott, America). For the intraperitoneal insulin tolerance test (IPITT) measurement, mice were fed overnight. After measurement of their postprandial blood glucose levels, the mice were intraperitoneally injected with 1 U/kg insulin. The subsequent measurement of blood glucose levels was similar with IPGTT.

### Cell lines

293T cells were purchased from the Cell Bank Type Culture Collection of the Chinese Academy of Sciences. It was cultured in DMEM supplemented with 10% fetal bovine serum (FBS), 10 U/mL penicillin and 100 mg/mL streptomycin. The cells were cultured in a humidified CO_2_ incubator at 37°C.

AML12 was purchased from the Cell Bank Type Culture Collection of the Chinese Academy of Sciences. AML12 was cultured in DMEM F12 supplemented with 10% FBS, 0.005 mg/ml insulin, 0.005 mg/ml transferrin, 5 ng/ml selenium, 40 ng/ml dexamethasone, 10 U/mL penicillin and 100 mg/mL streptomycin in a humidified CO_2_ incubator at 37°C.

### Mdivi-1 treatment

The Drp1 GTPase activity inhibitor Mdivi-1 was purchased from Topscience, China (T1907) and dissolved in DMSO to create a stock concentration of 10 mg/ml. 50 mg/kg Mdivi-1 was given to mice by intraperitoneal injection every two days for 2 weeks. The vehicle group received the same quantity of DMSO.

### Exhaustive swimming assay

Eight-week old WT and Clock^Δ19^ mice were put in a swimming box with a water depth of 20 cm and temperature of 25℃. During this process, the mice were forced to keep swimming by stirring the surrounding water. Mice were saved when they could no longer keep their nose above the water and started inhaling. Their swimming times were then recorded.

### Isolation of primary hepatocyte and liver mitochondria

Primary hepatocytes were isolated by perfusion of D-hanks and collagenase Ⅳ through the postcava to the portal vein. The liver was transferred to DMEM after perfusion, disintegrated by tweezers and filtered through a 70-μm cell strainer (BD Bioscience). Then, 90% percent percoll solution was used to separate the activated hepatocytes from the dead ones. The isolated hepatocytes were then cultured in DMEM with 10% FBS with the addition of penicillin-streptomycin solution.

Liver mitochondria were isolated in MSHE buffer (70 mM sucrose, 210 mM mannitol, 5 mM HEPES, 1 mM EGTA, and 0.5% fatty acid free BSA. pH 7.2). A piece of liver tissue was first homogenized in ~10-fold volume of cold MSHE buffer and centrifuged at 800 g for 10 min to remove the tissue fragments. The supernatant was centrifuged again at 8000 g for 10 min to obtain the crude mitochondria. The pallet was then resuspended in 100 μl of MSHE buffer, and the protein quantities were measured by BCA kit.

### Electron and confocal microscopy

Mitochondria number and diameter of liver and primary hepatocytes were analyzed by electron microscope. To visually study the mitochondrial dynamics, primary hepatocytes were transfected with Ad-cox8a-GFP/RFP for 24 hours and then photographed under a confocal microscope.

### Mitochondrial respiration assessment

For isolated mitochondria coupling and electron flow analysis, an XF24 Seahorse analyzer was used. Twenty micrograms of isolated mitochondria were plated on an assay plate in the initiation buffer (70 mM sucrose, 220 mM mannitol, 10 mM KH_2_PO_4_, 5 mM MgCl_2_, 2 mM HEPES, 1.0 mM EGTA and 0.2% fatty acid -free BSA, pH 7.2) plus substrates. For the coupling assay,10 mM succinate and 2 μM rotenone were used. For the electron flow assay, 10 mM pyruvate, 2 mM malate and 4 μM FCCP were used. The plate was then centrifuged at 2000 g for 20 min at 4℃. After centrifugation, the mitochondria were viewed briefly under the microscope and transfer to a 37℃ incubator for 10 min to warm. The plate was then transferred to the analyzer, and the experiment was initiated. The injections were as follows: for the coupling assay, 40 mM ADP, 25 μg/ml oligomycin, 40 μM FCCP and 40 μM antimycin were injected, while for the electron flow assay, 20 μM rotenone, 100 mM succinate, 40 μM antimycin and 100 mM ascorbate plus 1 mM TMPD were injected.

For the primary hepatocyte coupling assay, 25,000 cells/well were plated on a plate in advance in DMEM overnight to allow attachment. On the day of the experiment, the culture media was exchanged for basal medium (Sigma, D5030) with the addition of 10 mM pyruvate. The injections were as follows: 5 mg/ml oligomycin, 5 mM FCCP and 5 mM rotenone.

### Assessment of mitochondrial membrane potential, ROS production

WT and Clock^Δ19^ primary hepatocytes were cultured in DMEM overnight to adhere and stained with a specific reagent. To measure the mitochondrial membrane potential, after the removal of the culture medium and washing with PBS, the cells were stained with 5 μg/ml JC-1 (Beyotime Biotechnology, China, C2005) at 37℃ for 30 min. Then, they were washed with PBS 3 times before imaging. Similarly, the total cellular ROS were traced by 10 μM DCFH-DA fluorescence probe (Beyotime Biotechnology, China, S0033).

### Luciferase reporter assay

For the luciferase reporter assay, mouse Mrps24, Mrpl50 promoter and the mitochondrial D-loop were cloned into pGL3-Basic vector and fused with firefly luciferase. 293T cells were seeded and grown to ~80% confluency overnight and then transfected with the fusion plasmids with or without Clock and Bmal1 (3 replicates). After 24 h of transfection, the cells were harvested, and the luciferase activity was determined with a luminometer.

### Gene and protein expression measurement

Real-time PCR and western blot were used to measure the relative expression of mitochondrial dynamic genes and other genes. The primers used are listed in Table S1. The antibodies used are listed in Table S2.

### RNA immunoprecipitation

First, 10^7^ Aml12 cells (or a piece of liver) were crosslinked with UV, harvested in cold PBS and resuspended in 1 ml of RIP buffer [150 mM KCl, 25 mM Tris pH 7.4, 5 mM EDTA, 5 mM DTT, 5% NP40, 1 mM PMSF, 100 U/ml SUPERase· In™ RNase Inhibitor (Thermo Fisher)]. The cells (liver tissue) were then mechanically sheared using a Dounce homogenizer with 15–20 strokes (with a tissue homogenate) and briefly sonicated (OFF:30 s; ON:30 s for 13 mins). After incubation on ice for 10 min, the lysate was centrifuged at 12000 rpm for 10 min and split into three fractions of 45 μl each and two of 450 μl each (for Input, Mock and IP). Three micrograms of anti-CLOCK (Abcam, ab3517), or anti-IgG (Abcam, ab172730) antibody was added to the supernatant and incubated for 4 hours at 4°C with gentle rotation. Then, 40 μl of protein A/G beads were added and incubated for another 2 hours at 4°C with gentle rotation. The beads were pelleted at 2,500 rpm for 30 s. The supernatant was removed, and the beads were washed with RIP buffer 5 times. The beads were then resuspended with 100 μl of RIP buffer, and 10% were analyzed by Western blot with CLOCK-specific antibody. The remaining 90% were then digested with proteinase K and DNase Ι. The coprecipitated RNAs were further isolated by Trizol reagent. RT-PCR was then performed with specific primers.

### Statistical analysis

Data are presented as the means ± SEM. Statistical comparisons were conducted with unpaired Student’s t-tests or one-way ANOVA, as appropriate, and p < 0.05 was considered statistically significant.

## Acknowledgements

We thank professor Chi-h-Hao Lee for his kind providing of Cox8-RFP and Cox8-GFP adenovirus. We thank Dr Changpo Lin, Jieyu Guo and Mengping Jia for their helpful discussions and sharing of reagents. This work was supported by the National Science Foundation Fostering Talents in Basic Research of China No. J1310009 (RZ.Q.); the National Natural Science Foundation of China (NSFC) No. 81570771 (RZ.Q.), No. 81322003 (N.S.), No. 31571527 (N.S.), the Recruitment Program of Global Experts of the Organization Department of the Central Committee of the CPC (N.S.); the Science and Technology Commission of Shanghai Municipality (No. 13JC1401704) (N.S.).

## Author contributions

Conceptualization, Qian R and Lu C; Methodology, Lu C, Xu L, Li X, Sun N, Yan Z; Investigation, Xu L, Cheng Q, Hua B, Cai T, Lin J, Yuan G; Writing – Original Draft, Xu L; Writing – Review & Editing, Xu L, Lu C and Qian R; Funding Acquisition, Qian R and Lu C; Supervision, Qian R and Lu C.

